# The Power of Two: integrating deep diffusion models and variational autoencoders for single-cell transcriptomics analysis

**DOI:** 10.1101/2023.04.13.536789

**Authors:** Mehrshad Sadria, Anita Layton

## Abstract

Discovering a lower-dimensional embedding of single-cell data can greatly improve downstream analysis. The embedding should encapsulate both the high-level semantics and low-level variations in order to be meaningful and interpretable. Although current generative models have been used to learn such a low-dimensional representation, they have several limitations. Here, we introduce scVAEDer, a scalable deep-learning model that combines the power of variational autoencoders and deep diffusion models to learn a meaningful representation which can capture both global semantics and local variations in the data. By using the learned embedding, we show that scVAEDer can generate novel scRNA-seq data, predict the effect of the perturbation on various cell types, identify changes in gene expression during dedifferentiation, and detect master regulators in a biological process.

## Introduction

Recent advancements in single-cell sequencing technologies have presented an unparalleled opportunity to examine the characterization of cell types and states with unprecedented resolution and scale. Owing to this technology we can now generate vast amounts of data to explore intracellular and intercellular interactions and precisely quantify genomic, transcriptomic, and other multi-omics information at the cellular level (1). single-cell RNA sequencing (scRNA-seq) can also be used in conjunction with other technologies such as CRISPR-based perturbation (Perturb-seq) to uncover the function of genes and regulatory elements (2). Despite its potential, the technical complexity and high implementation costs of Perturb-seq have limited its widespread adoption (3).

Although scRNA-seq datasets have high dimensional nature, their intrinsic dimensionality is often low, owing to the cell type-specific gene regulatory network (GRN) (4). Machine learning (ML) methods have shown promising results in discovering these GRNs, enabling researchers to identify previously unknown cell types and characterize complex cellular processes during development and diseases (5,6). Among the many widely-used methods for analyzing single-cell data, Deep learning (DL) models, including autoencoders (AEs), are widely used, in different tasks such as dimensionality reduction, clustering, data denoising, and batch effect correction (7–9).

In addition, there has been a surge of interest in using deep generative models, such as variational autoencoders (VAEs) and generative adversarial networks (GANs), to analyze scRNA-seq data (10,11). By leveraging their ability to learn the underlying data distribution, these models have become valuable tools for generating realistic simulations of gene expression profiles, reconstructing cellular trajectories, and predicting cellular responses to different types of perturbations (12,13). While these generative models have demonstrated their potential, there are still issues that need to be addressed. Specifically, VAEs can experience posterior collapse (14), and GANs can struggle mode collapse (15), in which the learned mapping reduces significant portions of the input data to a single point in the output. Such problems are often caused by the disparity between the geometries of the prior and target data distributions (16). The performance of these models is also affected by other issues such as training instability, vanishing gradients, and prior hole (17). All these issues can eventually lead to biased or low-quality results, ultimately hindering the application and efficiency of using generative models in single-cell technology (18,19).

Recently, Denoising Diffusion Models (DDMs) have gained significant attention as potent generative models. They show their exceptional performance surpassing other generative models not only in image synthesis but also in domains such as natural language processing, music, and protein structure generation (20–22). In DDMs, the data undergoes a gradual perturbation through the diffusion process, then a deep neural network is trained to learn and remove the added noise in multiple steps. These methods can subsequently be employed to generate novel data in an iterative manner, starting from random noise without having issues associated with VAEs and GANs that were previously mentioned (23). However, traditional DDMs necessitate a costly, iterative sampling process and lack a low-dimensional latent representation (24). They employ a Markov process to transform the data distribution through small perturbations, which demands a large number of diffusion steps in both the training and inference phases. As a result, employing DDMs for large datasets is often not practical or feasible. Therefore, to enhance the quality and speed of these models, various attempts have been made. These include refining the forward process, implementing a more effective sampling method, integrating guidance from a classifier, or utilizing the low dimensional latent space of autoencoders for training (24–27).

In this study, we present scVAEDer, a new method for generating high-quality scRNA-seq data by combining the strengths of VAEs and DDMs. Instead of relying solely on VAE’s prior, our method incorporates both VAE and DDM priors to more precisely capture the distribution of latent encodings in the data. This allows for effective estimation of intermediate cell states and precise prediction of gene expression changes during the transition process from one cell type to another.

Furthermore, by using vector arithmetic in the DDM space scVAEDer outperforms state-of-the-art (SOTA) methods in predicting single-cell perturbation responses for novel conditions, which are not included in the training process. Finally, we showed that our method can successfully identify known and novel master regulatory genes involved in cellular reprogramming by computing the velocity of genes during interpolation. We evaluated the performance of our method by applying it to multiple datasets with diverse sizes and characteristics.

## Results

### Method outline

Generative models have shown great potential in analyzing single-cell data usually by finding a lower dimension which facilitates downstream tasks such as visualization, clustering, and data generation. However, they are not without their limitations. For example, in VAEs one significant issue is the prior hole problem, where the distribution of all encodings of the training data does not perfectly form a Gaussian due to a mismatch between approximate posterior and prior distributions. This can negatively affect the quality of the generated data when interpolating or operating directly on the VAE’s latent space.

We develop scVAEDer that combines the strengths of VAEs and DDMs to produce faster, higher quality, and more reliable results without encountering VAE issues. First, it takes single-cell gene expression data as input to train a VAE. Then it uses the latent embedding generated by the VAE to train the DDM which tries to model the real data distribution by learning a denoising process iteratively. Once trained successfully, the reverse process of diffusion can be used as a generative model which maps an arbitrary Gaussian noise *N*(*0, I*) to a new VAE latent representation after T successive steps. Therefore, by sampling from the trained DDM and feeding it to the VAE’s decoder, realistic gene expression profiles for different cell types can be generated. scVAEDer’s ability to generate high-quality data is due to its access to both VAE and DDM latent spaces, which enables capturing high-level semantics and low-level stochastic variations together. Using the trained embedding, we can predict the effects of different perturbations and perform interpolations between cell types in two different ways (Method section for more details). We apply scVAEDer to different biological data and show that the method can accurately approximate the distribution and trend of key genes during perturbation or cellular transition better than SOTA generative models. Also, as biological systems need to respond quickly to external and internal perturbations, we believe master regulators can be discovered based on their rate of response. scVAEDer computes the velocities by looking at the changes during interpolation steps or considering the average velocity over N steps. Gene Set Enrichment Analysis (GSEA) on genes that are ranked based on high velocity reveals enrichment in several crucial pathways.

### scVAEDer generates novel realistic scRNA-seq data

The application of DDMs in computer vision has yielded promising results for generating high-quality images. With this in mind, we investigate the potential of these models in generating single-cell gene expression data. By using scRNA-seq data of zebrafish hematopoiesis *x*_*0*_ with 1390 cells and 1845 genes, we train a VAE and compute the latent code *Z*_*sem*_ = Encoder(x_*0*_). Then, *Z*_*sem*_ is used to train a diffusion model. Figure 2a illustrates the input data is transformed into Gaussian noise through a gradual perturbation process using forward diffusion. Once DDM is trained, the reverse process can be utilized to generate new data through an iterative denoising process, beginning with random Gaussian noise. Increasing the number of steps in the training of DDMs results in higher-quality data generation, but also increases computational costs presenting a tradeoff between sample quality and cost. We generate new samples using the fully trained diffusion model and evaluate their quality. Our results show that scVAEDer generates high-quality data with excellent fidelity to the original one, exhibiting visual coherence and structural consistency that is especially critical for scRNA-seq data (Fig. 2a). Furthermore, we demonstrate that the quality of data synthesis achieved by the diffusion model is superior to the ones generated by sampling from VAE’s prior. Figure 2d provides clear evidence that the structure of the embedding is lost when just using the VAE prior, while the structure of the embedding generated by the diffusion model (Fig. 2c) is similar to the real data embedding (Fig. 2b). In addition, we compute the Total Variation Distance (TVD) between the real data embedding and those generated by sampling from VAE and scVAEDer to quantitatively determine which prior can generate samples closer to the true embedding. The samples produced by scVAEDer are significantly closer to the actual values and more precisely capture the distribution of latent embedding of data in comparison to the VAE model (Fig. 2e).

### scVAEDer models the transition between cells

The ability of cells to dedifferentiate is crucial in regenerative medicine and disease treatments. Leveraging scVAEDer’s access to the entire low-dimensional embedding, we seek to investigate gene expression changes by interpolating between different cell types in the latent space using hematopoiesis data. Specifically, we aim to predict how the expression of key genes changes as we interpolate between monocytes to Hematopoietic Stem/Progenitor Cells (HSPC). To avoid the “prior hole” problem in VAEs, we interpolate in the prior space of the latent DDM. Therefore, we first compute the latent embedding for monocytes and HSPCs using the VAE encoder and use DDM forward pass to perfectly map them to the Gaussian prior (Figure 3a). As there are no prior holes in that space, we can safely move along the interpolation path using 2000 equidistant steps between monocytes and HSPCs, and then use the reverse process to achieve the interpolation results in VAE’s latent space. These results can then be fed into the VAE’s decoder to understand changes in expression patterns as we move along the interpolation trajectory in the gene space. Interestingly, there are no samples generated in the empty region during the transition, which suggests that scVAEDer learns the correct complex dynamic of dedifferentiation (red dots in Fig. 3b).

Additionally, we compute the correlation between the cells generated by interpolation and the average gene expression of monocytes and HSPCs. During the initial interpolation stages, the cells exhibit greater similarity to monocytes; but as the process proceeds, they become more similar to HSPCs (Fig. 3c). Furthermore, by investigating the expression pattern of key genes, we can gain a deeper understanding of the fundamental mechanisms driving cell dedifferentiation. As can be seen in Figure 3d, there is a notable increase in the expression of HSPC markers in zebrafish, such as txnipa, sox4a, itga2b, and banf1 (28–31), which are critical regulators of stem cell maintenance and self-renewal. There is also a decrease in the expression of monocyte markers, such as cebpb, npsn, lyz, and fcer1gl (32–35), which are essential for the maturation and function of monocytes (Supplementary Figure 1 includes the trend of more genes). This suggests that scVAEDer can be used to understand the shifts in the activity of key genes during dedifferentiation. Moreover, by adjusting the number of diffusion steps before the interpolation process, we can modify the level of detail in the generated cells (Supplementary Figure 2). This finding is in agreement with the results reported by Ho et al (21).

**Fig. 1.**
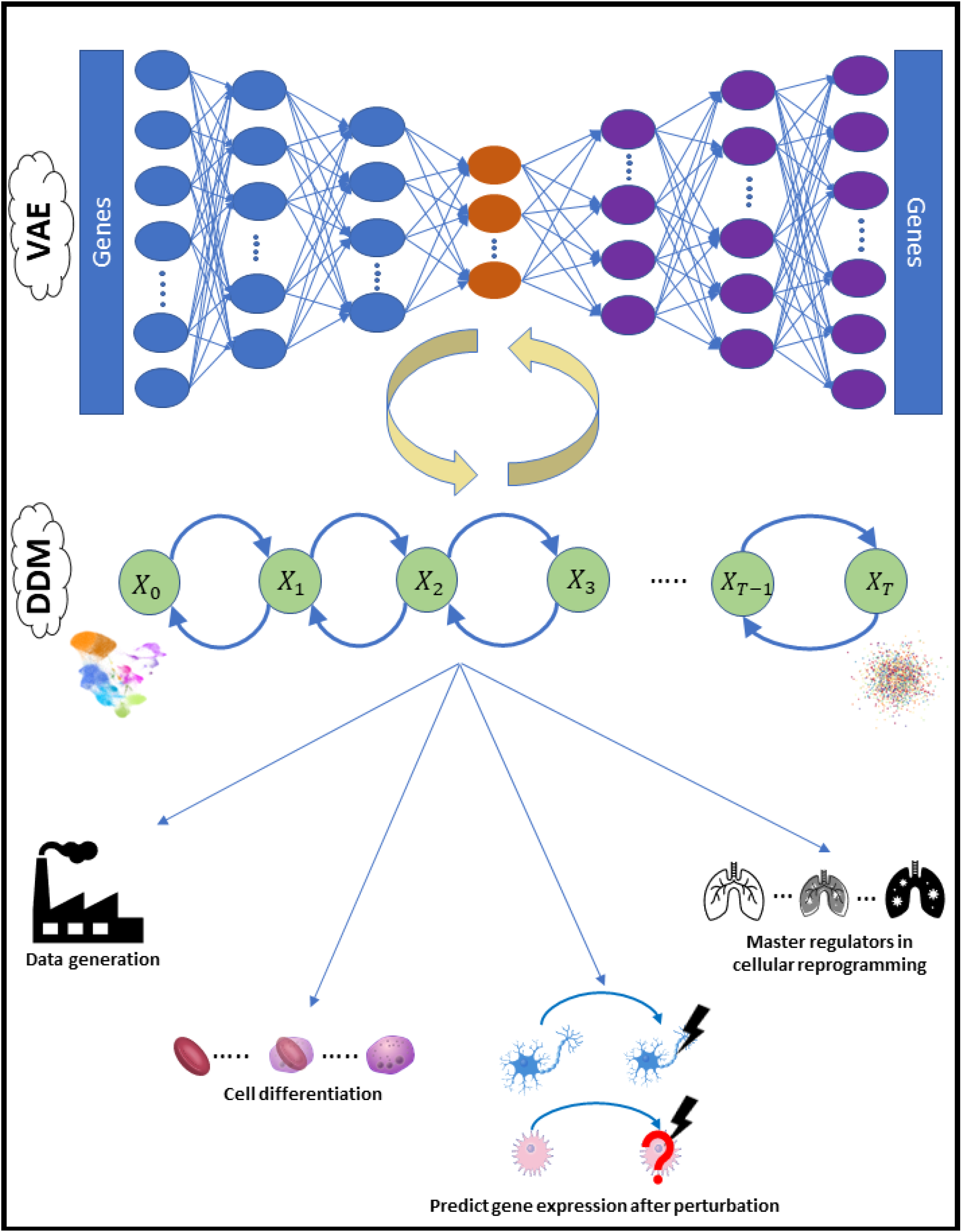
scVAEDer overview. scVAEDer integrates VAE and DDMs. First, a VAE is trained using the gene expression data. Then the VAE latent embedding is used to train the DDM. Combining together the model is able to decode back the gene space with high accuracy. scVAEDer can be used for different downstream analysis tasks such as generating high-quality single-cell data, understanding changes in gene expression during cellular differentiation, predicting the effect of perturbations on new cell types when expression data is available for multiple conditions, detecting master regulators by interpolating between different cellular states and ranking fast responder genes based on their velocity values.

**Fig. 2.**
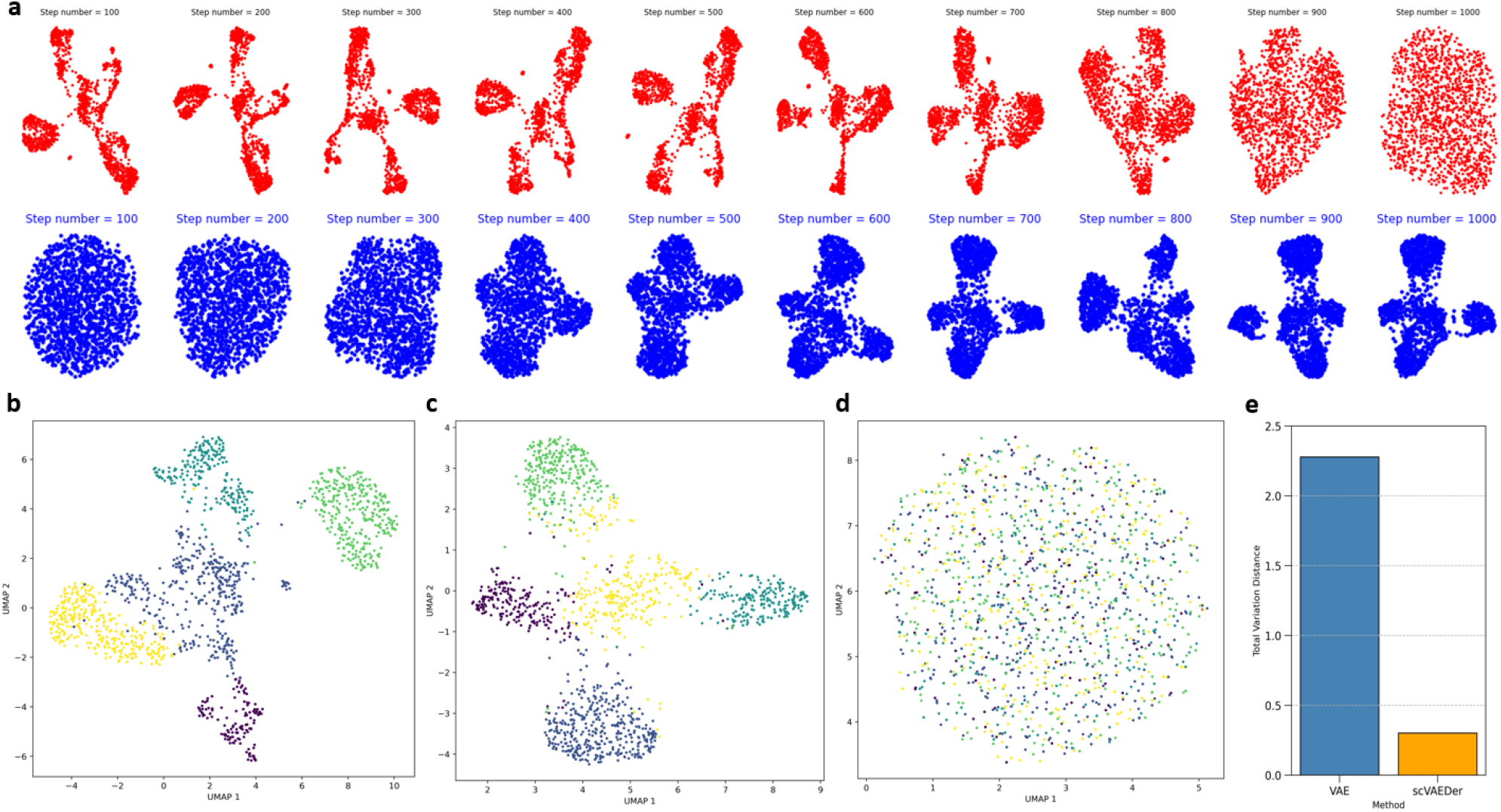
scVAEDer accurately learns the latent representation and generates new high-quality scRNA-seq data. a, red, forward Diffusion process with 1000 steps on hematopoiesis in zebrafish as we add noise to the data; blue, reverse process as the model learns how to denoise. b, UMAP visualization of the real data embedding. c, samples generated from DDM prior. d, samples generated from the VAE. e, Total Variation Distance between latent embedding of data and samples generated from the DDM as well as VAE prior distributions.

**Fig. 3.**
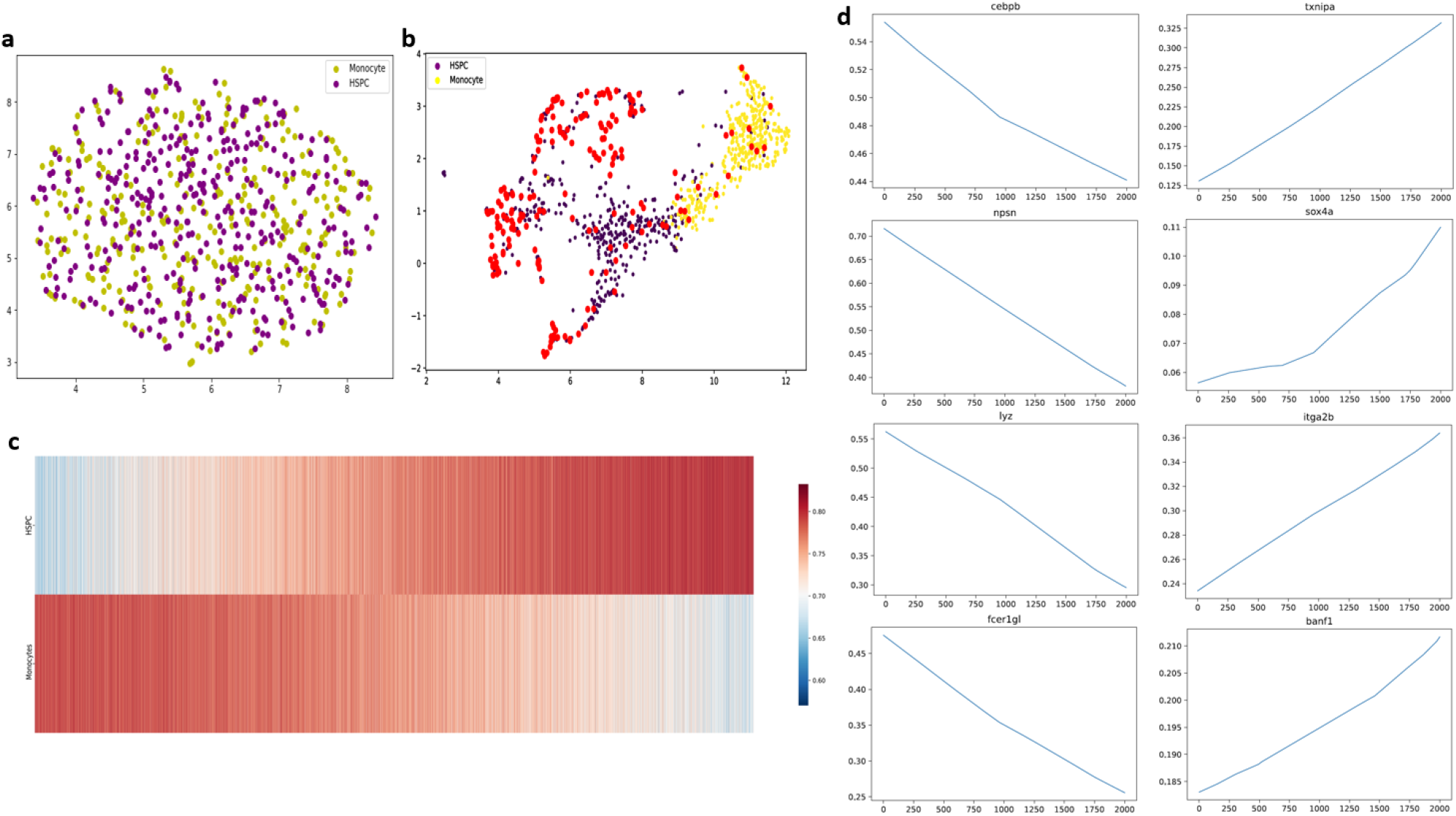
scVAEDer can be used to understand cellular dedifferentiation. a, Mapping HSPC and monocyte cells into the latent prior of DDM. b, Using DDM (1000 diffusion steps) and performing latent linear interpolation with 2000 equidistant samples (red dots). The absence of any sample generated in the empty region suggests that the model has learned an accurate embedding. c, Heatmap showing the similarity between gene expression of generated states and the real average expression of HSPCs and monocytes (using 100 DDM steps). d, Expression of selected marker genes along the interpolation path.

### scVAEDer predicts perturbation responses using feature manipulation

By studying cell responses to specific perturbations, we can gain insights into underlying mechanisms of a biological process, such as gene regulation and signal transduction, as well as the mechanisms of diseases. AEs are often used in the exploration of the effect of various types of perturbations on cells through vector arithmetic (5,11). scVAEDer latent can be used to predict the refined perturbed gene expression while avoiding VAE issues. To evaluate the performance of our model in predicting perturbation response, scRNA-seq data of human peripheral blood mononuclear cells (PBMCs) stimulated with interferon (IFN-β) was utilized to train scVAEDer (Fig. 4a). This dataset includes 6359 control and 7217 stimulated cells, with dramatic changes in the transcriptional profiles of immune cells induced by IFN-β stimulation. scVAEDer successfully predicts the gene expression of stimulated T-cells that are not part of the training set and demonstrates a strong correlation between the predicted mean expression and real data across all genes (Fig. 4b). We also conduct additional experiments to assess the versatility of our method. By excluding stimulated cells of each cell type, we train new models and showed the stimulation prediction for each cell type. Our method is robust and could accurately predict the responses of all other cell types (Fig. 4b).

**Fig. 4.**
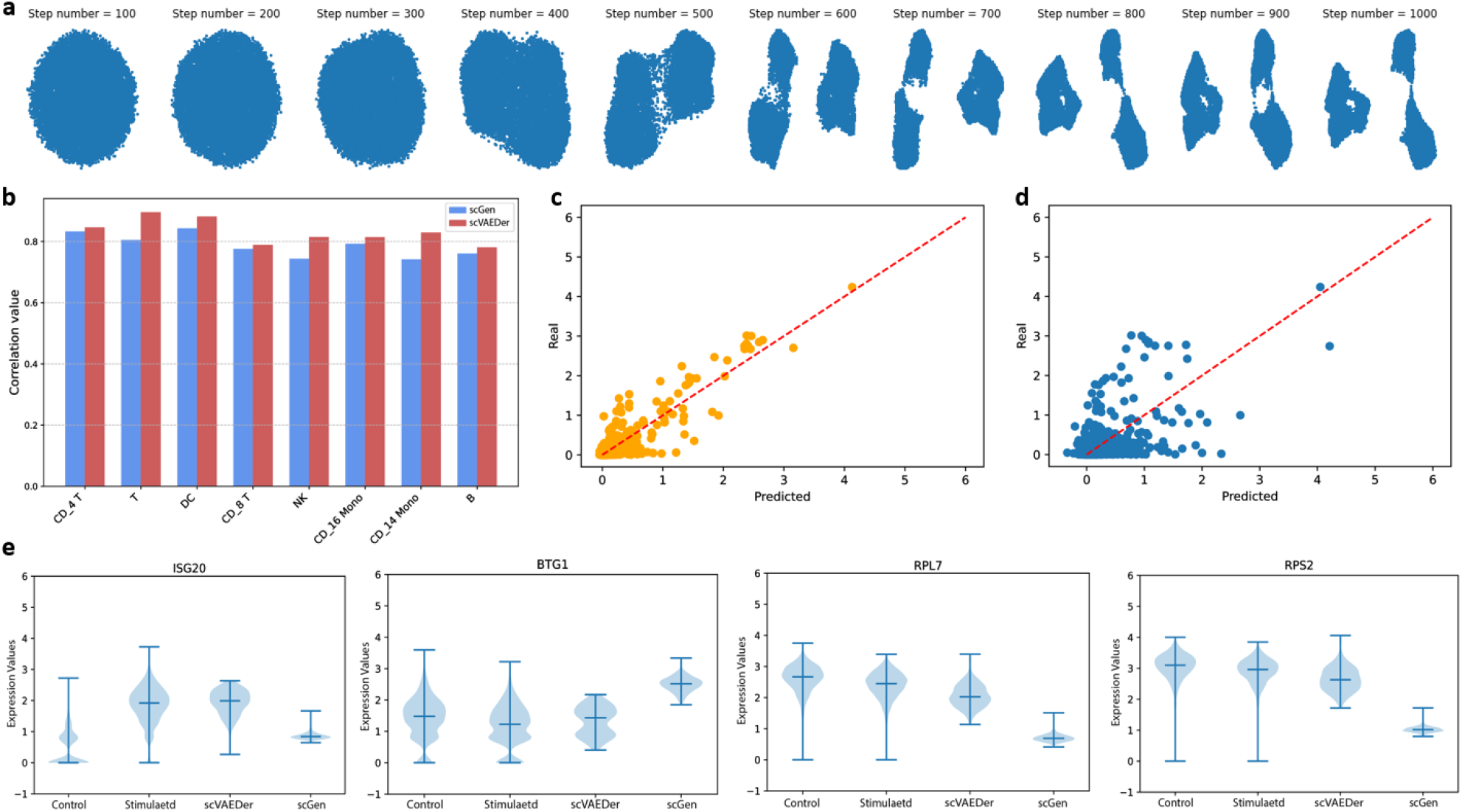
scVAEDer is more accurate than the SOTA method in predicting perturbation responses. a, Data generation using latent prior of DDM with 1000 diffusion steps. b, Comparison of the correlation values of average gene expression between real and predicted cells of various cell types obtained using scGen vs scVAEDer. c,d, Comparison between average gene expression of all genes between predicted and real stimulated T-cells using scVAEDer (c) and scGen (d). e, Violin plots for selected key genes across control, real stimulated, and stimulation predicted by scVAEDer and scGen.

We also conduct a comparison between scVAEDer and the state-of-the-art method scGen (which is based on VAE) by evaluating the correlation values between the predicted average gene expression and real data for T-cell across all genes (Fig. 4c,d respectively). We extend this analysis to other cell types and showed scVAEDer outperforms scGEN in predicting the effects of perturbation across different cell types (Fig. 4b). Furthermore, we use scVAEDer to predict the impact of the perturbation on the distribution of crucial genes and compare the results with scGen. scVAEDer achieves higher accuracy than scGen in predicting the gene distributions (Fig. 4e and Supplementary Figure 4 for more key genes).

### Disease progression and master regulators

The study of specific genes known as “master regulators” is of paramount importance in the control of cell fate and differentiation, as they can be targeted to reprogram cells into desired cell types and states. As such, their detection is crucial for gaining insights into disease mechanisms and advancing the development of potential therapies. Here we investigate the changes in gene expression during the transition from failed to reprogrammed cell states and to identify master regulators involved in this process. To achieve this, we employ an approach that involves interpolating between failed and reprogrammed cell states. Reprogramming data which has 18803 cells and 28001 genes is used to train scVAEDer after the preprocessing steps. We first confirm that the diffusion model is able to accurately reproduce the data from random Gaussian noise (Fig. 5a). Then using the trained model, we obtain the latent embeddings of reprogrammed and failed cells. The UMAP visualization of the cell states in the VAE space can be seen in Figure 5b. We use the average gene expression of failed and reprogrammed cells and interpolate in scVAEDer’s diffusion prior. Then the reverse process maps the interpolation results back into VAE’s space (Fig. 5c). The correlation values are computed between each interpolation step and the average gene expression of reprogrammed and failed cells. As we move along the interpolation path, the gene expression becomes more similar to the reprogrammed state and less like the failed state (Fig. 5d). We are also able to control the level of granularity during the interpolation process by adjusting the number of steps in the forward and reverse processes of DDM. Additionally, to understand marker genes during this transitional process, we calculate the absolute difference between the gene expression at time t=0 and t=3000 (corresponds to 20% of the total 15,000 as it corresponds to the global structural changes in the data). The genes with the highest absolute scores are displayed in Figure. 5e. This allows us to identify both known and previously unknown gene markers linked to the reprogramming of fibroblasts.

**Fig. 5.**
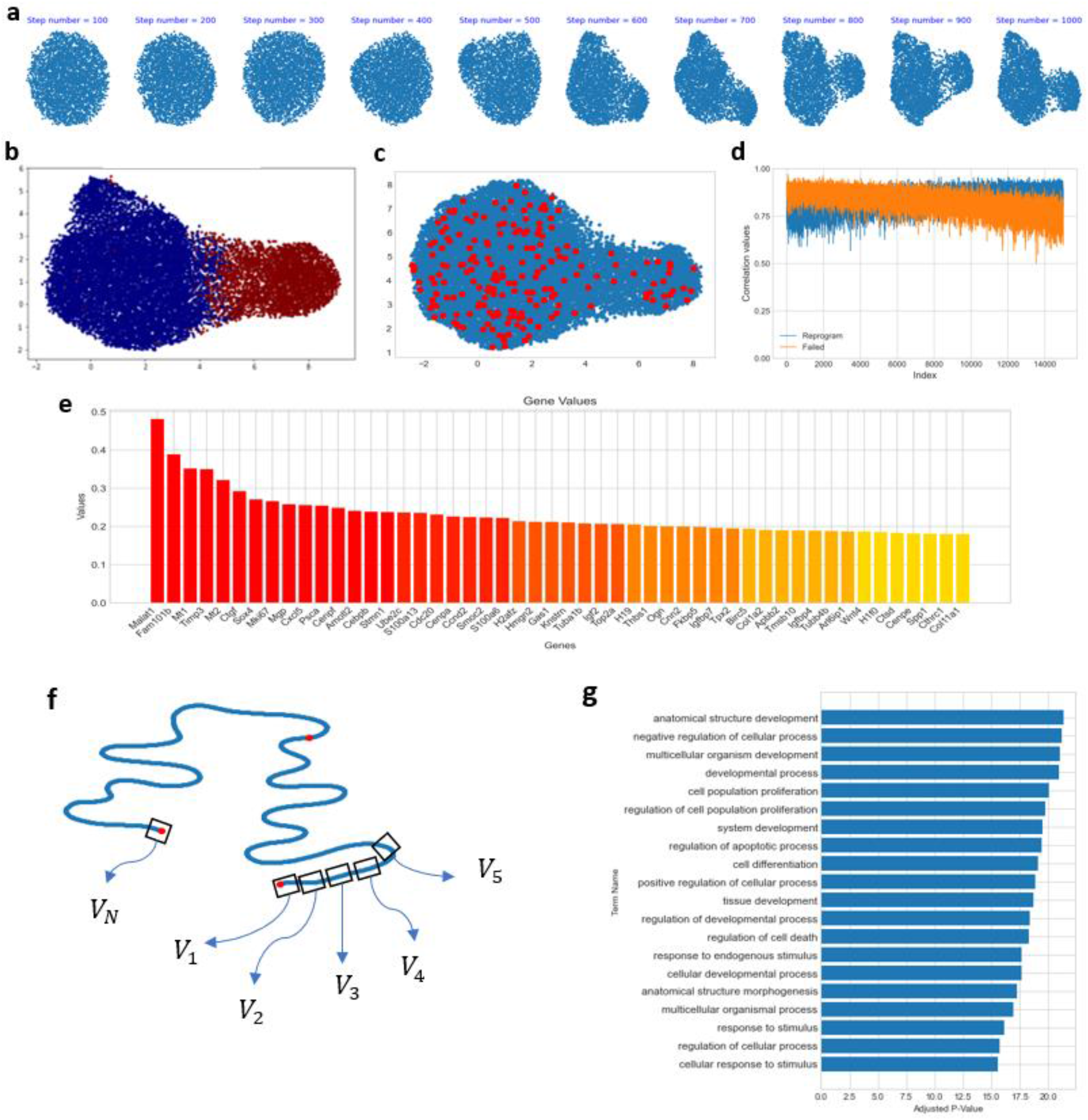
scVAEDer accurately detects master regulators during cellular reprogramming. a, The reverse process of DDM allows the generation of high-quality reprogramming samples from random Gaussian noise (the quality of generated samples is critical for downstream analysis). b, UMAP visualization of the data latent embedding, colored based on their state (red: failed, blue: reprogrammed). c, Data generated by interpolating between reprogrammed and failed states, represented by red dots. Remarkably, none of the interpolated samples are found outside the real representation of data. d, Correlation values between the new interpolated samples and the average gene expression of reprogrammed and failed cells, which demonstrates the shift in gene expression between failed and reprogrammed states. e) Ranking of genes based on high expression differences between t=0 and t=3000. f) Cartoon showing the process of computing gene velocities along the interpolation path to detect master regulators. g) Gene set enrichment analysis using 400 genes with the highest velocities, which reveals pathways that are crucial during the cell reprogramming process.

Next, we use scVAEDer to detect the master regulators involved in this process, by computing the gene velocities, given by the rate of changes in gene expression during interpolation. We believe gene velocities are informative as they allow us to identify crucial genes that should undergo rapid changes in order to play their central role in a specific biological process. By ranking genes based on their absolute velocity values, we detect key genes in this process, including Wnt4, Ptma, and FoxA1, which have been shown to be important in reprogramming and regeneration processes (the top 15 genes are listed in Supplementary Table 3). As a biological system needs different genes and pathways to be activated in order to succeed in different parts of the biological process, our approach makes it possible to detect master regulators in each step. Furthermore, we perform gene set enrichment analysis by selecting the top 10 genes with the highest velocities at each step of the interpolation process, with a focus on the first 40 steps (400 genes in total). Pathways related to morphogenesis, developmental process, and differentiation are significantly enriched, which demonstrates the ability of scVAEDer to detect related master regulators and pathways of biological processes.

## Discussion

Analyzing scRNA-seq data is a challenging task which involves identifying the appropriate underlying data embedding. The accuracy of this low-dimensional embedding is crucial for many downstream tasks, including cell clustering, cell type identification, gene expression analysis, and visualization. By learning the correct representation of data, we are also able to generate novel realistic data, predict the effect of different perturbations on cells, and detect the master regulators of a GRN or marker genes of a biological process. To produce high-quality samples, the generative model should also be able to take into account and control both high-level and low-level variations between different cell types.

Here we propose scVAEDer, a novel method that unifies the power of VAE and DDM. scVAEDer can generate realistic gene expression data using the latent representations of VAE for high-level and DDM for low-level variations. scVAEDer has access to the lower dimension of data which enables us to predict the effect of different unseen perturbations on a variety of cell types without having the limitations of the other generative models. We demonstrate using different datasets that scVAEDer outperforms the SOTA model in predicting the effect of perturbations. Additionally, the DDM prior of our method can be used for interpolation between different cell types, allowing us to generate new cells along the differentiation trajectory and identify highly expressed genes in each step. We also demonstrate that scVAEDer’s ability to compute gene velocity along the interpolation path can be useful in cellular reprogramming. Previous studies have demonstrated that master regulators are not typically the genes with the highest differential expression (36). To address this, we focus on genes with high velocities instead and we are able to discover both novel and previously identified master regulators and pathways that have been shown to play a vital role in cell fate and cellular reprogramming. In future versions of scVAEDer, it would be interesting to consider the diffusion model as a continuous process by using a stochastic diffusion equation (SDE) or a deterministic approach like probability flow ordinary differential equations (ODE).

## Methods

### Variational autoencoder (VAE)

VAE is a popular generative model that is widely used in many deep learning applications. It can generate new samples by capturing the underlying data distribution. It is based on the idea of maximizing a lower bound of the log-likelihood of the data. This is achieved by minimizing the KL divergence between the learned distribution and the true distribution. VAE models typically consist of two parts: an encoder that transfers input data to a latent space and a decoder network that maps latent space back to the original data space. The objective function can be written as:

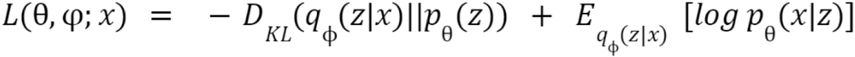

where *θ* and *ϕ* are the parameters of the decoder and encoder networks, *x* is the input data, *z* is the latent variable, *q*_ϕ_(*z*|*x*) is the approximate posterior distribution over the latent variables, and *p*_θ_(x|*z*) is the conditional distribution over the input data given the latent variables. In the equation above the first term is the KL divergence between the approximate posterior and the prior distribution. That encourages the learned distribution to be as close as possible to the prior distribution. The second term is reconstruction loss, which measures how well the model can reconstruct the input data from the latent embedding. By minimizing this objective function, the VAE model can learn to generate new samples similar to the input data.

### Deep Diffusion Model (DDM)

Diffusion models belong to a category of latent-variable models that have gained popularity in recent years. They can generate novel data from complex distributions, with higher quality compared to the existing generative models. To capture the underlying data distribution, DDMs follow a two-step process: a forward noising process and a reverse denoising process, working together in a coordinated manner. The forward process gradually destroys the structure of the data by adding noise, while the reverse process learns to recover the original data from the Gaussian noisy input.

It should be emphasized that the forward noising process is not trainable, but the reverse denoising process is parametrized and learned during training. In these processes, a first-order Markov chain with Gaussian transitions for the forward and backward processes is mostly used. The forward process can be written as:

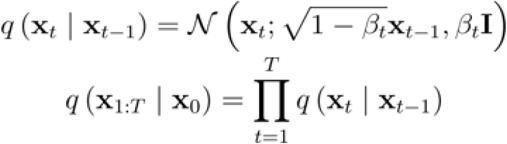

By choosing the correct values of the noise scheduler, we can assume *q*(*x*_*t*_| *x*_*0*_) is isotropic Gaussian. By using the <u>reparameterization trick</u>, we get:

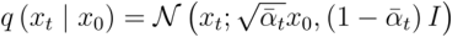

where:

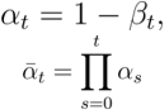

It’s worth mentioning that besides the Gaussian distribution, there are other distributions such as the gamma distribution that can be used. These distributions share a common property, which is the fact that the sum of two or more distributions results in the same distribution as well (37).

The reverse process can be formulated as:

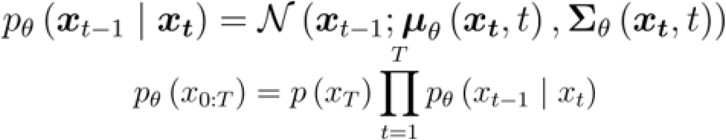

which starts from a random Gaussian X_*T*_: 𝒩 (X_*T*_, **0**,**I**) and in each step, the model learns. 𝒫 θ (*x*_0:*T*_)To train the reverse process, it has been shown (21) that the loss function below works well as it leads to a better result:

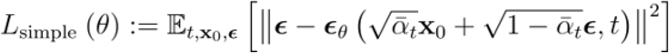

We can modify the above equation for our case as we train the diffusion model on the latent embedding of VAE. For the sake of simplicity, we ignore some of the weight terms (21). As a result, we express the DDM part of the scVAEDer objective as follows:

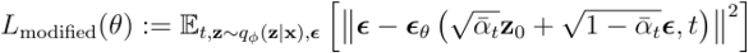

where *z* is the encoded value of *x* using the VAE’s encoder *q*_*ϕ*_ (*z*|x).

Once the training process is complete, it is possible to sample from the DDM prior iteratively using the sampling method below:

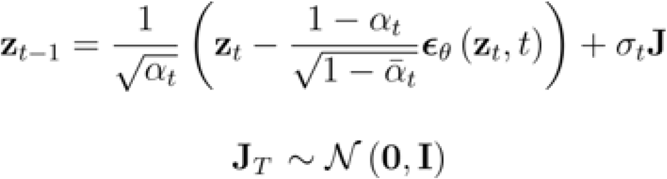

### Interpolation

A number of interpolation methods have been considered in previous generative models to understand the trajectory between two states. scVAEDer uses two distinct interpolation approaches:

1. By leveraging the VAE, we encode x_*1*_ and x_*2*_ (gene values) to obtain *z*_*1*_ and *z*_*2*_respectively in the VAE’s latent embedding. Afterward, by utilizing the diffusion forward process, we can obtain *q*_*1*_ and *q*_*2*_ in the prior space of DDMs. The interpolation can be performed in that space using, 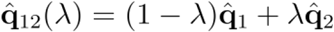 where λ is between [0,1]. Then the reverse process map 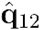 back to the VAE’s latent space and then by using the decoder of the VAE we can compute the interpolation values in the gene space.
2. Following a similar approach as the previous part, we compute *z*_*1*_ and *z*_*2*_ using the VAE’s encoder, and subsequently conduct interpolation using 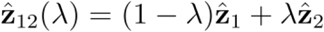. Then by adding Gaussian noise to 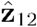, and performing the reverse diffusion process, we are able to generate better quality results than those produced solely by VAE. As in the previous section, the output of the reverse process is fed to the VAE’s decoder to map the embedding values from the VAE to the gene space.

To achieve more consistent results, different approaches can be utilized, such as an implicit instead of a probabilistic method (DDIM over DDPM) or the deterministic probability flow ODEs (22,38). Here we fix the stochasticity for all generated samples during interpolation to increase smoothness between samples. It is also essential to select the appropriate number of diffusion steps for each unique task, as this determines whether interpolation takes place at more refined or coarse levels of detail. When the diffusion step is zero, interpolation takes place in the VAE’s latent space. As we increase the number of diffusion steps, the initial interpolation information gradually diminishes, resulting in interpolations that are less detailed, coarser, and more varied (Supplementary Figure 2).

### Model architecture and training process

scVAEDer is trained on 4 Nvidia Tesla M60 machines with 32 GB GPUs, 448 GB system RAM, and 48 vCPUs. The training process involves two stages (detailed in Supplementary Tables. 1 and 2). The initial stage trains a VAE model, followed by the second stage where the diffusion model uses the latent embedding learned from the trained VAE. We chose a two-stage training process due to the higher memory and resource requirements of end-to-end training. Also, previous research has shown that end-to-end training can reduce both the quality of the generated data and the performance of the model (24). To ensure that the model can accurately learn the data embedding and generate high-quality samples, it is essential to select the correct parameters, such as the VAE and DDM network architectures, the number of diffusion steps, and the level of noise added during the forward process. Therefore, a hyperparameter search was performed on these parameters. Furthermore, an important aspect to consider when generating high-quality data is the number of steps and layers in the DDMs. Increasing these parameters can lead to better quality results, but the downside is that it requires significantly more computational resources. Therefore, it is crucial to consider the trade-off between the computational complexity and the quality of the results.

### Exponential Moving Average (EMA)

EMA can be used to improve the performance of the training process. Unlike the conventional methods which directly adjust model weights, EMA calculates a running average of the model’s weight parameters over the training process. This can lead to a more stable set of parameters. In our case, the weight parameters for the latent DDMs are tracked using EMA with a decay value of 0.999.

### Variance Scheduler

The noise schedules *β*_*1*_ to *β*_*T*_ are chosen such that the chain approximately converges to a standard Gaussian distribution after T steps. The variance parameter can be fixed to a constant or chosen as a schedule over the timesteps. To improve the performance, we incorporated three distinct variance schedulers, such as linear, quadratic, and sigmoidal. The user has the option to select any of these schedulers. The beta values are between *1e*^−*5*^ and *0.5e*^−*2*^ (Supplementary Figure 3).

### For visualization, reduced dimension data

Clusters were visualized using the Python package “UMAP”.

### Data preprocessing

The scRNA-seq gene expression data are centered and log-normalized. 2000 highly variable genes were chosen. We also performed batch normalization on the latent layer of VAE before training DDM as we found it improves the convergence speed and accuracy of the model.

### Comparison with scGen

We first downloaded the scGen python package from https://github.com/theislab/scgen and configure the relevant experimental environment as required. Also, to ensure a fair comparison, we selected the same neural network architecture for both scVAEDer and scGen.

## Data availability

All datasets analyzed in the current study are publicly available and can be downloaded from their public repository. The zebrafish hematopoiesis (39) raw data can be found under the accession number E-MTAB-5530 on ArrayExpress. The raw H.poly dataset (40) is available on the NCBI’s Gene Expression Omnibus under accession number GSE92332 and the preprocessed one can be downloaded from: https://github.com/theislab/scgen-reproducibility/blob/master/code/DataDownloader.py. Raw reprogramming data (41) is available from the Gene Expression Omnibus under accession number GSE99915. The preprocessed version can be downloaded from Cospar (42) package https://cospar.readthedocs.io/.

